# Estimating predator functional responses using the times between prey captures

**DOI:** 10.1101/2020.07.19.208686

**Authors:** Kyle E. Coblentz, John P. DeLong

**Affiliations:** School of Biological Sciences, University of Nebraska – Lincoln, Lincoln, NE, USA

**Keywords:** exponential distribution, feeding rates, foraging, individual variation, jumping spiders, Predator-prey interactions, predator/prey traits, time-to-event model

## Abstract

1. Predator functional responses, which describe how predator feeding rates change with prey densities, are a core component of predator-prey theory. Given their importance, ecologists have measured thousands of predator functional responses. However, most of these studies have used a single standard experimental method that is ill-suited to address many current, pressing questions regarding functional responses.
2. We derive a new experimental design and statistical analysis that quantifies the parameters of predator functional responses by using the time between a predator’s feeding events and can be used with individual predators requiring only one or a few trials. We examine the feasibility of this experimental method and analysis by using simulations to examine the ability of the statistical model to estimate the ‘true’ functional response parameters from simulated data. We also perform a proof-of-concept experiment estimating the functional responses of two individual jumping spiders feeding on midges.
3. Our simulations show that the statistical method is capable of reliably estimating functional response parameters under a wide range of parameter values and sample sizes. Our proof-of-concept experiment illustrates that the experimental design and statistical method provided reasonable estimates of functional response parameters and good fits to the data for individual jumping spiders using only a few trials per individual.
4. By virtue of the fewer number of trials required to measure a functional response, the method derived here promises to expand the questions that can be addressed using functional response experiments and the systems for which functional responses can be measured. For example, this method is well-poised to address questions such as intraspecific variation in predator functional response parameters and the role of predator and prey traits and abiotic conditions on shaping functional responses. We hope, therefore, that this time-between-captures method will refine our understanding of functional responses and thereby our understanding of predator-prey interactions more generally.

## Introduction

Predator functional responses are integral to ecological theory as they describe predator feeding rates given prey densities (Solomon, 1949; Holling, 1959b). Thus, functional responses are central to determining predator-prey interaction strengths that in turn can determine species coexistence (Paine, 1966; Holt, 1977; Coblentz & DeLong, 2020), the stability of ecological systems (Emmerson & Yearsley, 2004; McCann, 2011; Gellner & McCann, 2016), and many other important characteristics of predator-prey interactions and food webs. It is no surprise then that ecologists have spent decades performing experiments to measure functional responses. For example, in a recent effort to compile a database of functional response experiments (the Functional Responses from Around the Globe in all Ecosystems or FoRAGE database), Uiterwaal et al. (2018) collated over 2,000 functional response experiments from the literature. Despite this monumental effort devoted to measuring functional responses, there remain many open questions regarding functional responses that are difficult to address using the traditional ‘gold standard’ functional response experimental design.

Most functional response experiments follow the same experimental design. The experimenters choose a number of prey densities at which to measure foraging. The experimenters then add the specified number of prey to an arena generally containing a single predator individual and allow the predator to feed for a pre-specified amount of time that is generally constant across all prey densities. After the allotted foraging time, the experimenters record the number of prey killed. An alternative to this approach is similar but involves recording the amount of time it takes to forage to a specified number of prey (e.g. Gustafsson, Bergman, & Greenberg, 2010). This is repeated across all the prey densities multiple times typically using different predator individuals for each trial. The experimenters then fit a functional response model (or models) to the data using one of several existing statistical methods depending on whether prey were replaced as they were eaten (Royama, 1971; Rogers, 1972; Bolker, 2008; Rosenbaum & Rall, 2018; Uszko, Diehl, & Wickman, 2020).

Although the same basic experimental design has worked for thousands of functional response experiments, it is ill-equipped to address several open questions regarding functional responses. First, assessing the magnitude and ecological effects of intraspecific variation in predator functional response parameters is currently of great interest (Bolnick et al., 2011; Toscano & Griffen, 2014; Schröder, Kalinkat, & Arlinghaus, 2016). Using the current experimental design to measure an individual predator’s functional response at five prey densities with three replicates at each prey density would require fifteen trials with a single individual. Even this modest amount of replication and prey densities may quickly become infeasible for measuring the functional responses of tens of individuals. Furthermore, for rare predators or prey it may be extremely difficult to collect enough individuals to be able to perform a standard functional response experiment at all, limiting functional response experiments to abundant species or those that can be easily reared in a laboratory. Second, a key frontier in functional response experiments is understanding how predator and prey traits influence functional responses (Vucic-Pestic, Rall, Kalinkat, & Brose, 2010; Rall et al., 2012; Kalinkat et al., 2013; Uiterwaal & DeLong, 2020). The link between morphology and behavior to the functional response is integral to understanding selection on both predator and prey, but including several levels of predator and prey traits can quickly require a number of feeding trials that renders experiments infeasible. This problem only gets worse when realizing that foraging interactions are determined by the way predator and prey traits combine to influence strategies and the probability of successful attacks. For example, Kalinkat et al. (2013) examined the effects of predator and prey body sizes on the functional response across 25 different predator species feeding on eight differently sized prey species requiring 2,564 experimental units. Finally, foraging rates depend on both abiotic and biotic conditions (Abrams & Ginzburg, 2000; Gilbert et al., 2014; Preston et al., 2018; DeLong & Lyon, 2020), so identifying the way in which temperature, predator density, or habitat complexity, for example, influence the functional response generates the same level of replication challenge.

Below, we derive an alternative functional response experimental design and statistical analysis that uses a single predator individual and the time between its feeding events as it forages to measure its functional response. This method has the advantage of requiring only one or a few trials to estimate the entire functional response for an individual predator, easing some of the aforementioned difficulties with the current, widely used functional response experimental design. After deriving this method, we then use a simulation study to show that this method performs well at estimating functional response parameters when its statistical and biological assumptions are met under many circumstances. We then present a proof-of-concept example using Bold Jumping Spiders (*Phidippus audax*) foraging on midges (Chironomidae spp.).

### Derivation of the Experimental Method and Statistical Analysis

An intuitive motivation for our method of estimating functional responses is rooted in the idea of an individual predator ‘feeding down’ its functional response (Figure 1A). That is, as a predator feeds, they reduce the density of prey available. This fact was recognized long ago, and methods have been developed to account for prey depletion but not to capitalize on it (Royama, 1971; Rogers, 1972; Bolker, 2008). As a predator feeds down its functional response, the feeding rate should also change as the predator changes the prey density, and this relationship between feeding rate and density is exactly what a functional response is meant to capture. The question then becomes: Can we estimate the predator’s functional response from observations of the predator as it feeds on prey and depletes their number? The answer is yes, if we take advantage of the fact that the reciprocal of a rate is the expected time for an event to occur. Assume that the predator exhibits a Holling Type II functional response (as we will for the remainder of this manuscript; Holling, 1959a) in which the feeding rate, *f*, of the predator as a function of the resource density, *R*, is:

**Figure 1.**
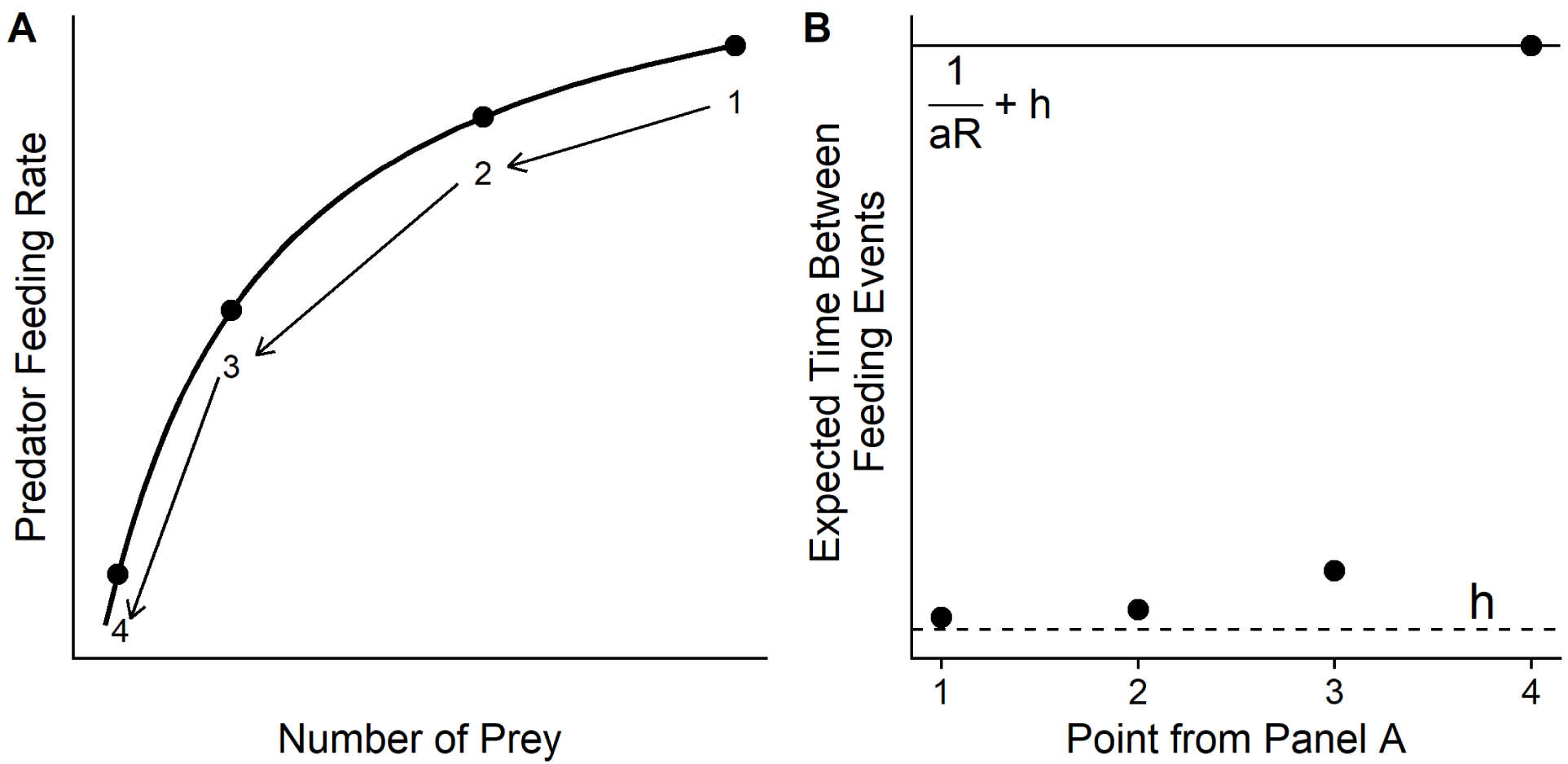
As predators ‘feed down’ their functional response, the number of prey decreases along with the predator’s feeding rate according to the functional response (e.g. from points 1→ 2 → 3 → 4 in A). Likewise, at each number of prey available, the expected time between feeding events according to the functional response also changes in a manner that provides information on the functional response parameters (B). In particular, times between feeding events at high prey densities are particularly informative about the handling time (*h*, point 1), and times between feeding events at low prey densities are particularly informative about the space clearance rate given the handling time (*a*, point 4; *R* in panel B is the number of prey available).

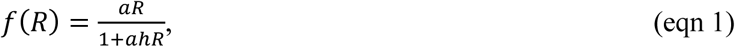

where *a* is space clearance rate of the predator (aka attack rate or attack efficiency) and *h* is the handling time. Under this model, the expected time between feeding events is the reciprocal of the feeding rate, or:

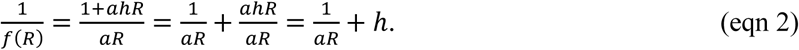

Therefore, at prey density, *R*, the expected time-to-feed for the predator is the expected time to encounter and catch a prey individual 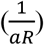 plus the time required to handle the previous prey (*h*). If we watch a predator as it feeds across different prey densities and record the time between its feeding events, we should then be able to use the time-to-feed data to infer the predator’s functional response parameters.

To derive a statistical method for estimating the functional response, we translate equation 2 into a stochastic process. First, we focus on the time to encounter and catch a prey individual 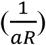. A widely used statistical distribution for time-to-event-data is the Exponential distribution, which models the time between events in Poisson processes given a constant rate of event occurrence, β. We can assume then that at prey density, *R*_*i*_, the time to encounter and catch a prey individual is Exponentially distributed with β = *aR*_*i*_, because this term describes the mass action encounters among predator and prey that lead to foraging events. To extend this model from the expected time to encounter and catch a prey individual to the expected time between feeding events, we also need to add the handling time. To do so, we recognize that the minimum time between feeding events is the handling time. Thus, we can alter the Exponential distribution so that its minimum value is the handling time. Using this, we can model the time-to-capture, *y*_*i*_, for a predator at prey density, *R*_*i*_, as:

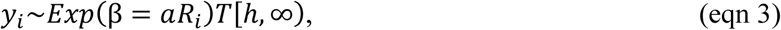

where the tilde (∼) means ‘distributed as’, Exp denotes the Exponential distribution, and T denotes that the distribution is truncated to the interval [*h*, ∞). With a series of time-to-capture measurements across a range of prey densities, one can use equation 3 to estimate a and h from the data using Bayesian or maximum likelihood methods (Bayesian methods are employed throughout the manuscript, but a maximum likelihood approach is outlined in Online Supplementary Material 1).

To build intuition on how the time between feeding events can allow us to infer the values of the space clearance rate and handling time, we consider two examples. First, assume that the prey density is very large. At large prey densities the time to encounter and capture a prey item 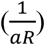 is very small. Thus, the time between feeding events is approximately equal to the handling time (Figure 1B). Alternatively, assume that there is only one prey item. The expected time-to-feed for the predator is 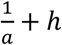 (Figure 1B). So, given the handling time, the time-to-capture allows us to directly infer the space clearance rate. Therefore, by following the times between feeding events as a predator ‘feeds down’ its functional response, we can estimate the functional response parameters. Furthermore, observations at high prey densities are particularly informative about the predator’s handling time, and observations at low prey density are particularly informative about the predator’s space clearance rate (we will return to this later in discussing alternative experimental designs).

Although our derivation of this alternative experiment for measuring predator functional responses comes directly from the definition of the functional response, it is not a given that the statistical method will accurately estimate the functional response parameter values or that this method is logistically feasible given predator behavior in the real world. To address this, we first performed a simulation study in which we varied the functional response parameters, simulated time-to-capture data sets from the statistical model and assessed how well the model was able to estimate the true functional response parameter values. We also performed a proof-of-concept experiment using Bold Jumping Spiders feeding on midges to illustrate and evaluate the use of this alternative experimental approach to estimating functional responses. Overall, the simulations and experiment suggest that the experimental approach outlined here is feasible for many organisms. We hope that adopting this approach can help to expand the kinds of questions that can be addressed using functional response experiments and thereby increase our understanding of predator-prey interactions and their consequences.

## Materials and Methods

### Simulation Study

We used simulations to examine the ability of the statistical model to estimate functional response parameters from simulated time-to-capture data. For three values of space clearance rates (a = 0.003, 0.5, 50), three values of handling times (h = 0.001, 0.01, 1), and three values of initial prey densities (10, 25, 50), we simulated the times between feeding events until all prey were consumed. At each prey density, the time-to-capture was simulated by drawing a random sample from an exponential distribution with rate β = α*R* and adding the handling time. After simulating the data, we then estimated the space clearance rates and handling times by fitting the model in equation 3 to the simulated data in a Bayesian framework using the program ‘Stan’ through the package ‘RStan’ in R (Carpenter et al., 2017; R Core Team, 2019; Stan Development Team, 2019). We placed a Cauchy prior with location μ = 0 and scale σ = 5 truncated at zero for the low and intermediate values of the space clearance rate and a Cauchy prior with μ =50 and σ = 15 truncated at zero for the highest value of the space clearance rate. For the handling time., we placed a Cauchy prior with μ = 0 and σ = 2.5 truncated below at zero and above at the minimum observed time between feeding events (the minimum possible time between feeding events provides an upper limit on the handling time). We chose these priors for the simulations to reflect the use of vaguely informative priors which we encourage given the large number of previous functional response experiments. We performed one hundred simulations for each space clearance rate, handling time, and initial prey density combination. After fitting the statistical models, we determined: 1) the proportion of simulations for which the true space clearance rates and handling times were within the 95% Credible Interval of the estimates, 2) the proportion of parameter point estimates that were greater than the true value (over-estimated), and 3) the mean absolute difference between the point parameter estimates and the true value. The range of parameters considered are well within those that have been observed in traditional functional response experiments (Uiterwaal et al., 2018). Some of the parameter combinations and initial prey densities used for the simulations led to total times required to consume all prey that would be difficult to achieve experimentally (alternative experimental designs may be possible for predators in which consuming all prey is unlikely or would take too much time; see Discussion). However, our goal with the simulations was only to examine the ability of the statistical method to estimate the functional response parameters over a range of values. Code to perform simulations is available so that researchers can assess the efficacy of our method for their own use (see Data Availability).

### Empirical Study – Bold Jumping Spiders

As a proof-of-concept experiment, we used the time-to-capture method derived here to estimate the functional responses of two Bold Jumping Spiders (*Phidippus audax*) feeding on small adult midges (Chironomidae spp., average length 61.5 mm, SD = 12). To gather the time between capture data, we used a camera (HERO3, GoPro, Inc., San Mateo, CA, USA) to record video of the spiders foraging in a circular clear plastic arena 25cm in diameter. For the first spider (Spider 1; 1.04 cm length including abdomen and cephalothorax), we performed three feeding trials on three consecutive days. We placed three prey in the arena in the first trial, eleven in the second, and eight in the third trial. We chose this sequence of prey levels intentionally to minimize satiation effects on subsequent days. Spider 1 consumed all of the prey in the first and third trials and eight of the eleven prey in the second trial. For the second spider (Spider 2; 1.07 cm length including abdomen and cephalothorax), we also performed three trials on three consecutive days. We placed three prey in the arena for the first and second trials and eleven prey in the arena for the third trial. Spider 2 consumed all of the prey in each of the trials. For each trial, we calculated the time between captures for each prey following the first capture, since by definition the time-to-capture for the first prey does not include handling time. Each spider had one observation for which it did not finish handling a prey before attacking the next one. As this is likely to bias the handling time estimates, we removed these observations prior to analysis. For the data from each spider separately and for the combined data across both spiders, we fit the exponential model in equation 3 in a Bayesian framework using the program Stan through the R package ‘RStan’ (Carpenter et al., 2017; Stan Development Team, 2019). We placed a vaguely informative Cauchy prior with μ = 10 and σ = 10 truncated at zero on the space clearance rate and a vaguely informative Cauchy prior with μ = 0 and σ = 2 truncated below at zero and above at the minimum observed time between feeding events. The priors were derived from the FoRAGE database using the observed space clearance rates and handling times for invertebrate predators feeding on invertebrate prey (Uiterwaal et al., 2018). A maximum likelihood approach to estimating the spider functional responses is outlined in Online Supplementary Material 1. All of the code and data to perform the analyses are available (see Data Availability).

## Results

### Simulation Study

Overall, the statistical model was able to estimate the functional response parameters well under most circumstances (Table 1). However, the model struggled to estimate handling times when both space clearance rates and handling times were low, generally exhibiting bad coverage of the Credible Intervals and overestimating handling times until reaching a space clearance rate of 0.5 and a handling time of 0.01 (Table 1, Figure 2A). As space clearance rates increased, handling times were more accurately estimated across all of the levels of handling times considered (Table 1, Figure 2B). The coverage of the 95% Credible Intervals for the space clearance rates were near their nominal values for nearly all the combinations of variables considered (Table 1). However, the models showed a tendency to overestimate space clearance rates when they were low (Table 1, Figure 2). Estimates of space clearance rates and handling times improved with higher sample sizes (Table 1). In general, these results support that the time-to-capture method is feasible under many realistic functional response parameter values.

**Table 1.** Summary of the ability of the introduced exponential statistical model to estimate space clearance rates and handling times using time-to-capture data under a range of parameter values and initial prey densities.

| Space Clearance Rate | Handling Time | Initial Prey Density | 95% Credible Interval Coverage of Space Clearance Rates | Mean Absolute Space Clearance Rate Difference from True Value | Percent Estimates Grater Than True Space Clearance Rate | 95% Credible Interval Coverage of Handling Times | Mean Absolute Handling Time Difference from True Value | Percent Estimates Greater Than True Handling Time |
| --- | --- | --- | --- | --- | --- | --- | --- | --- |
| 0.003 | 0.001 | 10 | 90% | 0.001 | 72% | 0% | 2.8 | 100% |
| 0.003 | 0.001 | 25 | 94% | 0.0005 | 61% | 5% | 0.7 | 99% |
| 0.003 | 0.001 | 50 | 94% | 0.0004 | 69% | 17% | 0.2 | 100% |
| 0.003 | 0.01 | 10 | 94% | 0.001 | 62% | 4% | 2.35 | 98% |
| 0.003 | 0.01 | 25 | 89% | 0.0007 | 67% | 17% | 0.7 | 92% |
| 0.003 | 0.01 | 50 | 96% | 0.0003 | 62% | 64% | 0.12 | 57% |
| 0.003 | 1 | 10 | 89% | 0.001 | 69% | 100% | 2.05 | 79% |
| 0.003 | 1 | 25 | 93% | 0.0005 | 62% | 95% | 0.6 | 68% |
| 0.003 | 1 | 50 | 95% | 0.0004 | 55% | 97% | 0.20 | 52% |
| 0.5 | 0.001 | 10 | 90% | 0.18 | 75% | 45% | 0.03 | 99% |
| 0.5 | 0.001 | 25 | 89% | 0.1 | 64% | 77% | 0.005 | 80% |
| 0.5 | 0.001 | 50 | 92% | 0.07 | 54% | 91% | 0.001 | 74% |
| 0.5 | 0.01 | 10 | 88% | 0.2 | 70% | 86% | 0.02 | 85% |
| 0.5 | 0.01 | 25 | 93% | 0.1 | 67% | 91% | 0.004 | 53% |
| 0.5 | 0.01 | 50 | 95% | 0.06 | 52% | 97% | 0.001 | 49% |
| 0.5 | 1 | 10 | 86% | 0.2 | 62% | 98% | 0.2 | 54% |
| 0.5 | 1 | 25 | 92% | 0.1 | 60% | 93% | 0.005 | 41% |
| 0.5 | 1 | 50 | 96% | 0.1 | 66% | 97% | 0.003 | 50% |
| 50 | 0.001 | 10 | 99% | 7.5 | 64% | 93% | 0.0003 | 47% |
| 50 | 0.001 | 25 | 95% | 6 | 63% | 97% | $3.5 \times 10^{-5}$ | 53% |
| 50 | 0.001 | 50 | 97% | 4.5 | 46% | 96% | $1 \times 10^{-5}$ | 53% |
| 50 | 0.01 | 10 | 99% | 6.7 | 54% | 98% | 0.0002 | 52% |
| 50 | 0.01 | 25 | 98% | 6.1 | 55% | 94% | $5 \times 10^{-5}$ | 58% |
| 50 | 0.01 | 50 | 96% | 4.6 | 48% | 95% | $1 \times 10^{-5}$ | 54% |
| 50 | 1 | 10 | 100% | 6.3 | 61% | 98% | $2.4 \times 10^{-4}$ | 48% |
| 50 | 1 | 25 | 99% | 4.9 | 58% | 92% | $4.5 \times 10^{-5}$ | 46% |
| 50 | 1 | 50 | 98% | 4.5 | 55% | 94% | $1 \times 10^{-5}$ | 48% |

**Figure 2.**
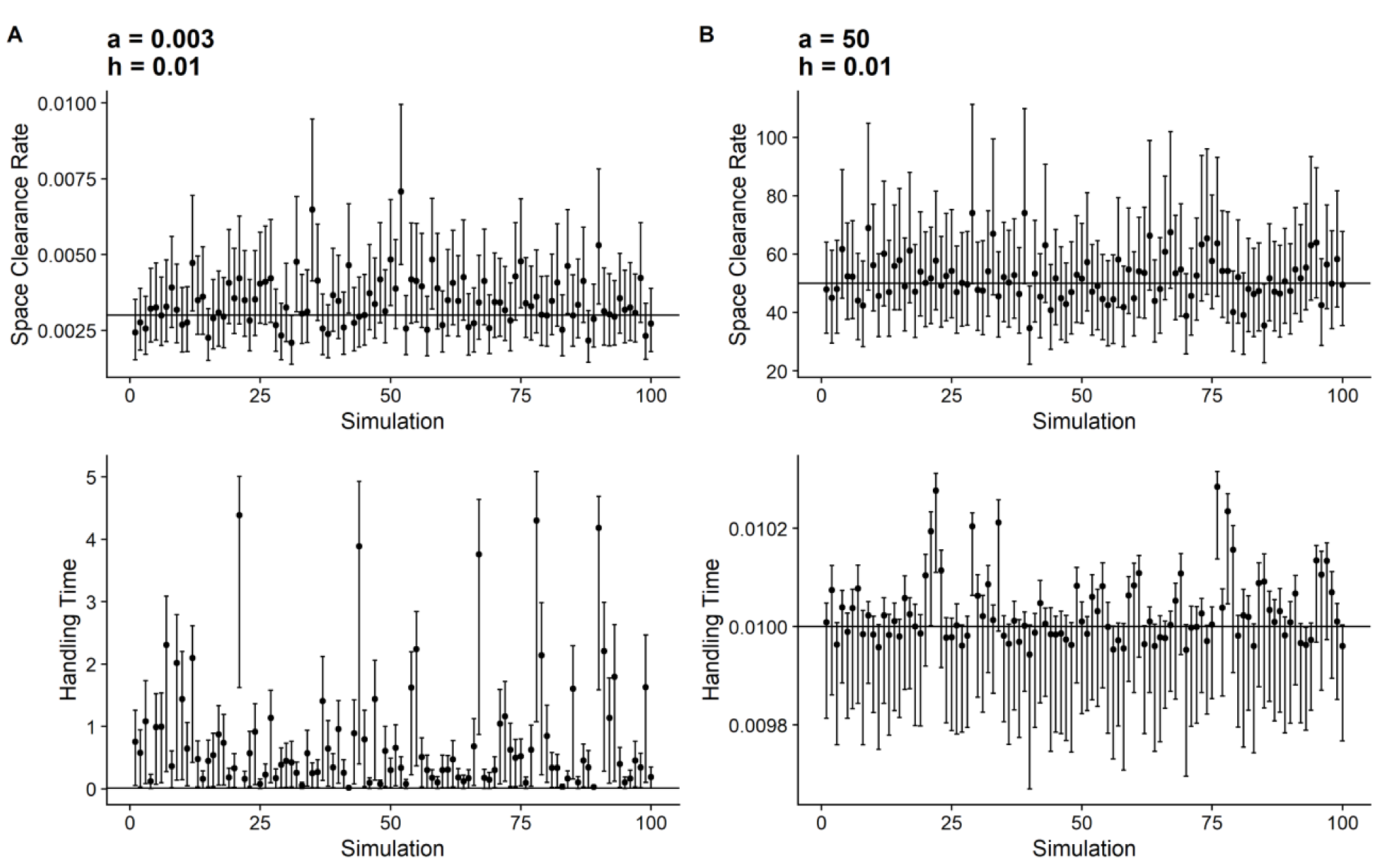
The analysis of simulated time-between-prey capture data shows that space clearance rates (*a*) are estimated well across a wide range of parameter values (A,B). The statistical method has difficulty estimating handling times (*h*) when space clearance rates are small (A), but estimates handling times well at larger space clearance rates (B). For a summary of the simulation results, see Table 1. These figures correspond to the results for the space clearance rates and handling times given in the figure and an initial prey density of 25.

### Empirical Study – Bold Jumping Spiders

For the time-to-capture data for both the individual spiders and the collated data across both individuals, the exponential model provided good fits to the data and reasonable estimates of space clearance rates and handling times (Figure 3, 4). We estimated Spider 1’s space clearance rate to be 6.5 m^2^ day^-1^ (95% Credible Interval (CrI): 3.6—10.9) and its handling time to be 0.00556 days (95% CrI: 0.004—0.0058). We estimated Spider 2’s space clearance rate to be 14.0 m^2^day^-1^ (95% CrI: 6.25—25.8) and its handling time to be 0.0035 days (95% CrI: 0.0024—0.0037). Using the data from both spiders, we estimated a functional response intermediate between the two individual functional responses with a space clearance rate of 7.2 m^2^day^-1^ (95% CrI: 4.6—10.4) and a handling time of 0.0035 days (95% CrI: 0.00279— 0.00367; Figure 4).

**Figure 3.**
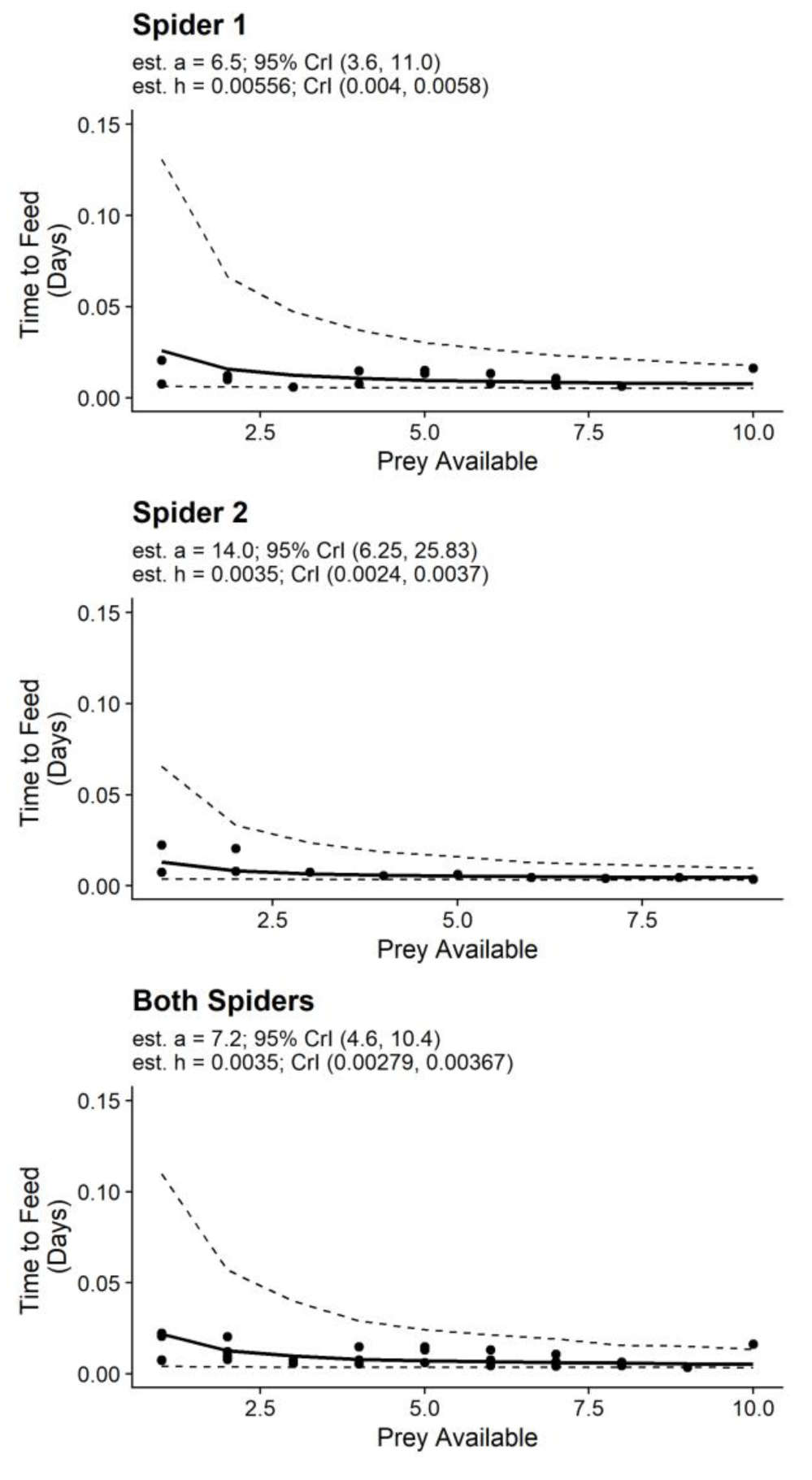
The statistical model provided good fits to the time between prey captures (Time to Feed) and the number of prey available and reasonable functional response parameter estimates. The solid line represents the expected relationship between the prey available and the time between prey captures using the point estimates of the functional response parameters. The dashed lines represent the 95% posterior predictive interval for the time between feeding events at each prey density.

**Figure 4.**
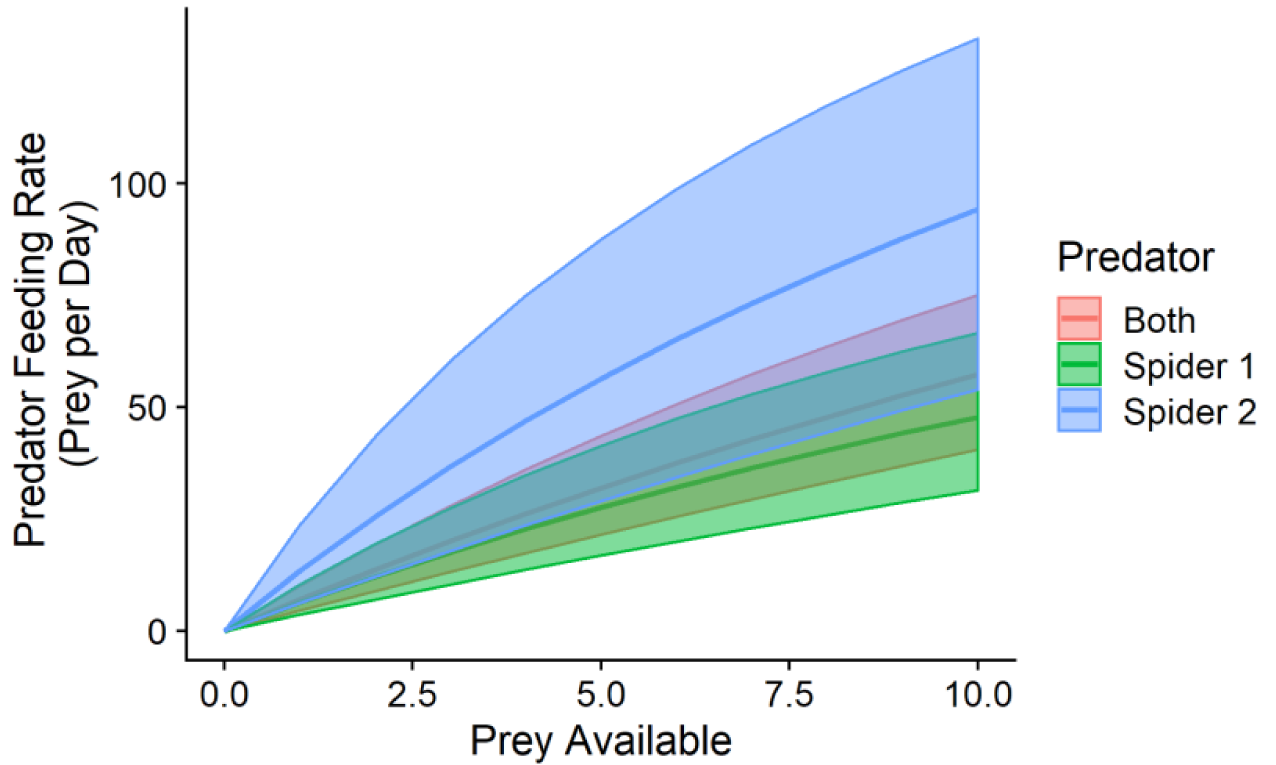
Predicted functional responses from individual jumping spiders (1 and 2) and the data from both spiders combined show generally higher feeding rates for Spider 2, lower feeding rates for Spider 1, and intermediate feeding rates for the combined data across both spiders. The colored areas represent 95% Credible Intervals for the functional responses.

## Discussion

Predator functional responses are an integral component of predator-prey theory and, thus, massive effort has been placed in measuring thousands of functional responses mainly using one traditional experimental design. This design has been effective, but it has also been limiting both in terms of the systems to which it can be applied and in its ability to help address pressing questions regarding functional responses such as intraspecific variability in functional response parameters, variability across prey types, and the role of traits in determining functional responses. We introduce an alternative method for estimating predator functional responses that uses the time between a predator’s prey captures to infer the parameters of its functional response. Because this method requires only one or a few trials to estimate an entire functional response (because each capture event is informative instead of just the capture total) and can be used with individual predators, we believe it has the potential to open several novel avenues of research on functional responses that would be difficult or impossible using current experimental methods for estimating predator functional responses.

Although the time-to-capture method has potential for furthering the study of functional responses, it is also no panacea. As with traditional functional responses experiments, the time-to-capture method will also be generally limited to species for which the experiments can be easily performed (i,e, species that can be placed in foraging arenas, aquaria, etc.). However, the time-to-capture method is less constrained in terms of predators and prey that are rare, as fewer individuals are necessary to measure a functional response. The simulation study also illustrates that the time-to-capture method is not likely to be able estimate handling times well for species that have small space clearance rates. This is because small space clearance rates lead to long times between prey capture events even at high prey densities, which can obscure the signal of handling times from the times between captures. The method also is unlikely to work for predators that continue to handle prey items as they attack and consume the next prey individual. As the handling time in the statistical model is the minimum amount of time between prey captures, the time-to-capture method is likely to underestimate handling times for predators that do not fully handle prey before capturing the next prey. Lastly, the time-to-capture method is likely to be most appropriate for actively foraging species under prey densities in which the predator does not become satiated during foraging given the density of prey available.

We derived the statistical method envisioning a predator ‘feeding down’ its functional response from some starting prey density, and we used this method to estimate the jumping spider functional responses. However, the statistical analysis only requires time between captures at various prey densities to estimate the functional response parameters. This creates the potential for alternative experimental designs and applications to non-experimental data. For example, for predators that consume prey slowly or become satiated quickly, one could focus on collecting the times between captures at very high and low prey densities with perhaps a few prey captures per trial. As previously mentioned, the times between prey captures at high and low prey densities are particularly informative about the predator handling times and space clearance rates, respectively. Thus, this design could be used to ‘lock down’ the estimates of these parameters while giving less attention to intermediate prey densities. Furthermore, additional trials at high or low prey densities may help with parameter estimation even if the ‘feeding down the functional response’ design is used. The statistical method here also may be used for observational data if the researcher is able to simultaneously observe prey capture events and the densities of prey. One intriguing application of this method to observational data is to carnivores or marine mammals outfitted with collars containing accelerometers. Previous studies have shown that these collars can identify predation events (Viviant, Monestiez, & Guinet, 2014; Wang et al., 2015). If one could also estimate prey densities at the locations of the predator, it may be possible to use the time between predation events to provide some estimate of the functional responses for predators that are not amenable to direct experimentation.

The application of the time-to-capture method to measure the jumping spider functional responses raises a broader point regarding the measurement of handling times in functional response experiments in general. The handling times estimated using the time-to-capture method while the jumping spiders were actively foraging are akin to the handling times in Holling’s original derivation of the Type II functional response or the Holling disc equation (Holling, 1959a). Holling derived the disc equation by assuming that all of a predator’s time was spent either searching for prey or handling prey. Thus, the handling time in the original disc equation was meant to represent the time unavailable for searching due to handling (but also includes digestion time if it takes away from search time; Jeschke, Kopp, & Tollrian, 2002). In contrast, in many functional response experiments, predators are placed in arenas for lengths of time that are likely to include not only active foraging by predators, but also inactivity, exploration unrelated to prey searching, and hygiene-related activities that will also decrease the time spent actively searching for prey. In these cases, all of the other non-handling activities that reduce predator searching time will contribute to the handling time. Thus, handling times estimated from functional responses are likely to depend on the lengths of trials and whether the predator is actively foraging for the entire length of the trial. This suggests that one should use caution directly comparing handling time estimates across functional response experiments regardless of the method used to estimate them. It is possible that by deriving a time budget for the predator one may be able to translate between different ‘handling time’ estimates.

## Conclusions

Here we have introduced a method for estimating predator functional responses using data on a predator’s time between prey captures. This method is a promising addition to existing methods for estimating predator functional responses such as traditional functional response experiments and observational methods for estimating functional responses in the field (Novak & Wootton, 2008; Novak, Wolf, Coblentz, & Shepard, 2017). Because this method only requires one or a few trials to estimate an entire functional response and can be performed on individuals, we believe that this method will expand the types of questions one can ask using functional response experiments and the systems in which functional responses can be measured. We also hope that future developments will extend this method to estimating alternative functional response forms beyond the Holling Type II response and will be modified to address questions such as predator dependence in functional responses, the effects of abiotic variables on functional response parameters, and the estimation of multispecies functional responses.

## Acknowledgements

This project was funded by a James S. McDonnell Foundation Grant to JPD.

## Authors’ Contributions

KEC and JPD conceived the ideas herein; KEC developed the statistical method, performed the simulation study, analyzed the functional response experiment data, and wrote the first draft of the manuscript; JPD performed the Bold Jumping Spider functional response experiment. All authors contributed to the final version of the manuscript and gave approval for publication.

## Data Availability

All data and code can be found at: https://github.com/KyleCoblentz/TimeToCaptureFRMethod. This GitHub repository will be permanently archived on Zenodo upon acceptance.

