## Supplementary material for "Estimating predator functional responses using the times between prey captures": Online Supplementary Material 1

Throughout the main text we use a Bayesian approach to fit the exponential model derived in the main text to the time between prey capture data. In recognition that researchers differ in their preferred statistical frameworks, here we outline a maximum likelihood approach to fitting the exponential model. We also illustrate this maximum likelihood approach by applying it to the same bold jumping spider data analyzed in the main text.

Again, our model for the observed times between prey captures, $y_i$, is

$$y_i \sim Exp(\beta = aR_i)T[h,\infty), \quad (1)$$

where $\sim$ stands for 'distributed as', $Exp$ denotes the exponential distribution, β is the rate parameter of the exponential distribution, $a$ is the predator's space clearance rate, $R_i$, is the prey density associated with the observation $y_i$, and $T[h,\infty)$ denotes that the distribution is truncated to the interval $[h,\infty)$.

Our goal is to develop a maximum likelihood approach to fit this model to the data and be able to get point estimates and confidence intervals for the functional response parameters $a$ and $h$. Unfortunately, doing so is not as straightforward as just applying standard maximum likelihood estimation software such as the 'bbmle' package in R (Bolker, 2008). The reason for this, is that the maximum likelihood estimate for the handling time lies on the boundary of its parameter space. When the maximum likelihood estimate of parameter lies on the boundary of its parameter space, standard tools for estimating parameters and their confidence intervals no longer apply (Bolker, 2008). To see that the maximum likelihood estimate for handling time lies on the boundary of its parameter space, consider the log likelihood ($\ell(a,h;y)$) of the above model,

$$\ell(a,h;y) = \sum_{i=1}^{n} log(aR_i) - aR_i(y_i - h), \quad (2)$$

where $n$ is the number of total observations, $log$ is the natural logarithm, and the remaining parameters are defined above. Now, also note that the values that $h$ can take are restricted between zero and the minimum time between prey captures or the minimum $y_i$, i.e. the shortest time between prey captures places an upper limit on how long handling a prey item can take. Now insert the minimum $y_i$ as the estimate for $h$ in equation 2. Our goal is to maximize the value of the log likelihood. When $h$ is equal to the minimum $y_i$, the right-most term in equation 2 becomes zero for the $y_i$ that is the minimum. Furthermore, for the other $y_i$'s the right-most term is reduced to the smallest possible value that still keeps $h$ within its defined boundaries. Therefore, the maximum likelihood estimate for $h$ is the minimum observed time between prey captures or the minimum $y_i$ and this estimate lies on the boundary of $h$'s parameter space.

Because we cannot directly apply standard maximum likelihood software to get estimates and confidence intervals for both $a$ and $h$, we instead take an iterative approach to getting the estimates. First, we calculate the maximum likelihood point estimate for $h$ which we have shown is the minimum observed time between prey captures. Next, we estimate $a$ and its confidence interval using the 'mle2' function in the R package 'bbmle' fixing the value of $h$ to its maximum likelihood estimate. We then place a lower bound on $h$ using a result from Epstein and Sobel (1954). Epstein and Sobel (1954) show that for a left-truncated exponential distribution (or a so-called two parameter exponential distribution) a lower confidence bound on the minimum of the distribution is given by,

$$y_1 - \frac{F_{2,2(n-1),1-\alpha}}{\beta n}, \qquad (3)$$

where $y_1$ is the minimum observed time between prey captures, $F_{2,2(n-1),1-\alpha}$ is the $1-\alpha$th quantile of an F distribution with 2 and 2(n-1) degrees of freedom, $n$ is the sample size, β is the rate parameter of the exponential distribution, and α is the desired level of confidence. Unfortunately, this result is for independent and identically distributed (iid) samples from a left-truncated exponential distribution. Note that in our case β is not a single value but is a function of $a$ and $R_i$. Thus, to get a value for β, we use the maximum likelihood point estimate for $a$, and mean value of the resource density, $R_i$, across the experiment. Below we use simulations to show that confidence intervals derived for the handling time this way exhibit good coverage of the true values but may be overly conservative. We furthermore demonstrate the use of this approach by applying it to the bold jumping spider data.

**Simulations**

As in the main text, we use simulations to examine the ability of the maximum likelihood approach outlined here to estimate the functional response parameters. As we have shown in the main text that the time-to-capture method has difficulty estimating handling times when space clearance rates are low, we focus on estimates of the functional response parameters when the space clearance rate is intermediate ($a = 0.5$) and handling times are intermediate or high ($h = 0.01$, 1) and for all of the handling times considered in the main text ($h = 0.001$. 0.01. and 1) when space clearance rates are high ($a = 50$). For each parameter combination, we also consider sample sizes of 10, 25, and 50 prey. At each combination of parameter values and sample size, we simulated 100 time-to-capture data sets from the exponential model. We then estimated the space clearance rates and handling times using the maximum likelihood procedure derived above. After fitting the statistical models, we determined: 1) the proportion of simulations for which the true space clearance rates and handling times were within the 95% Confidence Interval of the estimates, 2) the proportion of parameter point estimates that were greater than the true value (over-estimated), and 3) the mean absolute difference between the point parameter estimates and the true value. Code to perform the simulations and analyses are available (see Data Availability in the main text).

A summary of the results of the simulations are given in Table S1.1. Across all parameters and sample sizes, the maximum likelihood method provided good estimates for the space clearance rates and exhibited coverage of 95% confidence intervals near their nominal values. There was also a trend of slight bias of the space clearance rates estimates towards overestimating the space clearance rate. Because the maximum likelihood estimate for the handling time is the minimum observed time between prey captures, the handling time point estimate is always an overestimate of the 'true' handling time. The lower bound for the handling time appears to be conservative, exhibiting coverage of the confidence intervals often greater than the nominal 95%. Furthermore, when handling times are low and the sample size is low, the lower bound can be estimated to be negative. In these cases, the practitioner will have to settle for a point estimate of the handling time knowing that the true handling time is likely to be less than the point estimate or switch to the Bayesian implementation. That being said, the maximum likelihood estimate of the handling time was often quite accurate for the parameter values used.

**Bold Jumping Spider Analysis**

We analyzed the same Bold Jumping Spider data in the main text using the maximum likelihood approach derived above. The parameter estimates using the maximum likelihood method were comparable to those estimated using a Bayesian framework. For Spider 1, the maximum likelihood estimates for the space clearance rate and handling times were 7.9 $m^2 day^{-1}$ (95% Confidince Interval (CI) 4.6—12.7) and

0.00585 days (95% CI 0.0043--0.00585), respectively (the Bayesian estimates and 95% Credible Intervals were 6.5 (3.6—10.9) and 0.00556 (0.004—0.00585)). For Spider 2, the maximum likelihood estimates for the space clearance rate and handling times were 18.8 $m^2 day^{-1}$ (95% CI 9.8—32.3) and 0.0037 days (95% CI 0.0029--0.0037), respectively (the Bayesian estimates and intervals were 14 (6.25—25.8) and 0.0035 (0.0024—0.0037). The maximum likelihood estimates for the space clearance rate and handling time for the combined spider data were 8.0 $m^2 day^{-1}$ (95% CI 5.3—11.45) and 0.0037 days (95% CI 0.0023—0.0037), respectively (the Bayesian estimates and intervals were 7.2 (4.6—10.4) and 0.0035 (0.0028—0.0037). In general, the maximum likelihood approach estimated slightly higher space clearance rates due to the slightly higher handling time estimates compared to the Bayesian approach.

**Table S1.1:** Summary of the ability of the maximum likelihood approach to estimate space clearance rates and handling times from simulated time-to-capture data under a range of parameter values and initial prey densities.

| Space Clearance Rate | Handling Time | Initial Prey Density | 95% Confidence Interval Coverage of Space Clearance Rates | Mean Absolute Space Clearance Rate Difference from True Value | Percent Estimates Greater Than True Space Clearance Rate | 95% Confidence Interval Coverage of Handling Times | Mean Absolute Handling Time Difference from True Value | Percent Estimates Greater Than True Handling Time |
|---|---|---|---|---|---|---|---|---|
| 0.5 | 0.01 | 10 | 94% | 0.18 | 68% | 97% | 0.04 | 100% |
| 0.5 | 0.01 | 25 | 96% | 0.1 | 60% | 95% | 0.006 | 100% |
| 0.5 | 0.01 | 50 | 93% | 0.07 | 61% | 95% | 0.002 | 100% |
| 0.5 | 1 | 10 | 92% | 0.19 | 72% | 97% | 0.036 | 100% |
| 0.5 | 1 | 25 | 95% | 0.09 | 58% | 99% | 0.006 | 100% |
| 0.5 | 1 | 50 | 95% | 0.05 | 54% | 97% | 0.0015 | 100% |
| 50 | 0.001 | 10 | 92% | 20 | 65% | 98% | $3x10^{-4}$ | 100% |
| 50 | 0.001 | 25 | 95% | 8.9 | 55% | 99% | $6x10^{-5}$ | 100% |
| 50 | 0.001 | 50 | 93% | 6.3 | 57% | 95% | $1x10^{-5}$ | 100% |
| 50 | 0.01 | 10 | 96% | 15 | 59% | 98% | $4x10^{-4}$ | 100% |
| 50 | 0.01 | 25 | 98% | 10 | 66% | 98% | $6.6x10^{-5}$ | 100% |
| 50 | 0.01 | 50 | 97% | 5.5 | 52% | 97% | $1.5x10^{-5}$ | 100% |
| 50 | 1 | 10 | 93% | 18 | 66% | 95% | $3.6x10^{-4}$ | 100% |
| 50 | 1 | 25 | 95% | 8.8 | 53% | 99% | $6.6x10^{-5}$ | 100% |
| 50 | 1 | 50 | 97% | 5.9 | 53% | 95% | $1.7x10^{-5}$ | 100% |
